# Consensus Gene Expression Subtypes Of Epithelial Tumors

**DOI:** 10.1101/042234

**Authors:** Anguraj Sadanandam

## Abstract

While the heterogeneity of cancer is increasingly appreciated, it is also clear that recurrently deregulated pathways are shared between tumors across tissue types. To identify similarities in subsets of tumors from multiple epithelial organs, we performed transcriptome profile analysis of different epithelial tumors and cell lines (n=2928) from 17 different organs. We identified three to seven consensus gene expression subtypes in epithelial tumors and cell lines, irrespective of the organ-specific origin. These consensus subtypes showed biological significance. Moreover, we identified specific gene signatures that could be used to develop clinical assays for future application of this classification system. Overall, our study identifies consensus transcriptome subtypes across organ-specific tumors and highlights new avenues for precision therapy irrespective of the organ of origin.

## Introduction

The clinical management of solid tumors is currently organized and administered by tissue of origin and histopathology. Many studies have recently reported transcriptional subtypes in cancers arising from specific tissues (1-6). However, many clinically actionable aberrations cross tissue type; *KRAS* mutations in lung cancer are one example (7), and *ERBB2* amplification in breast cancer (8) is another. Recently, we published gene expression subtypes in pancreatic adenocarcinoma (PDA) (2), colorectal cancer (CRC) (3), and breast cancer (BrCA) (9) and observed similarities in gene expression in certain subtypes across these three tissues of origin. This led us to examine the similarities in subtypes across different epithelial organ type cancers based on publically available gene expression profiles. It is our expectation that the results from this study will help not only to understand similar mechanisms operating in subsets of cancers from different epithelial organs, but will also assist in the development of new methods to target cancer independent of the organ from which they originate.

## Results

### Identification of Consensus Subtypes

We obtained published gene expression microarray analysis of epithelial tumors (n=843) from five different organs – breast, colorectum, lung, pancreas and ovary (10-14), named as multiorgan tumor (MOT)-5 dataset. An unsupervised clustering analysis of MOT-5 data set with 353 genes using consensus based non-matrix factorization (NMF) algorithm (2, 3, 15) identified three tumor subtypes (Figure 1A) score. These subtypes were named as subtype-1, −2 and −3 (Figure 1A). Interestingly, we found that the MOT-5 samples can be further subdivided to form a total of seven subtypes (Figure 1A).

**Figure 1.**
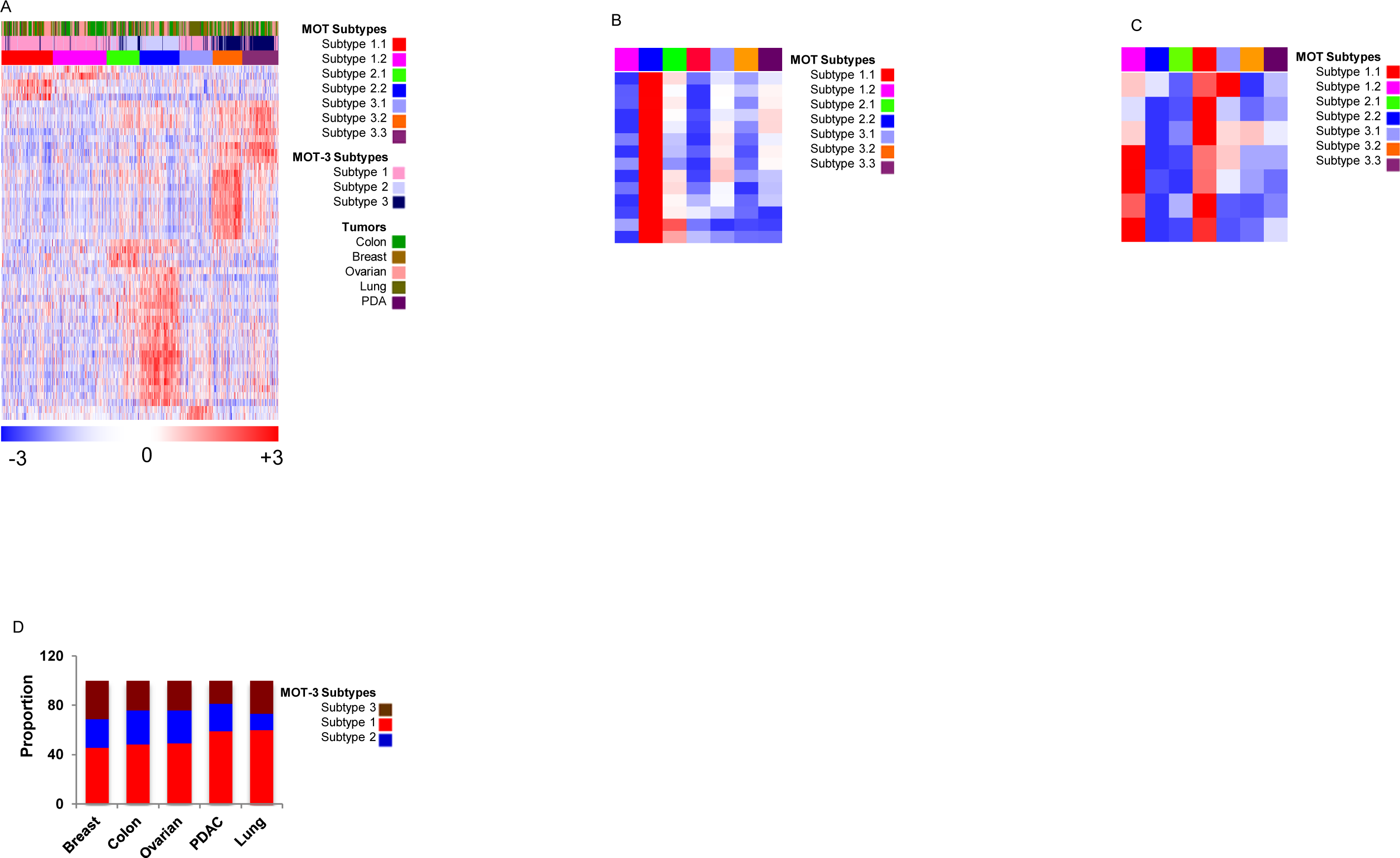
MOT subtypes and their signatures. **A**. Heatmap showing three and seven-subtype possibilities and their specific gene signatures (MOTassigner-51) using MOT samples. **B-C**. Heatmap with specific gene signature of **B**. subtype 2.1 and **C**. subtype 1.1. **D**. Proportion of three subtypes in different organ-type tumors.

### Characteristics of the MOT-5 Subtypes

Next, we performed significant analysis of microarrays (SAM) using the MOT-5 data set and identified 275 subtype-specific genes (MOTassigner-275) that are statistically significant (*FDR=0*). We further reduced the MOTassigner-275 gene signature to a 51 gene signature (MOTassigner-51), with the expectation that this may be useful for developing assays in the clinic (Figure 1A). The characteristic genes associated with each subtype were shown in Figures 1B-C.The proportions of subtypes across cancer types are shown in Figure 1D.

## Discussion

Personalized/precision medicine is revolutionizing the way cancers are treated by identifying and exploiting specific biological mechanisms/processes operating within a given tumor. We have attempted to understand the similarities across multiple epithelial organ-type cancers by subtyping them using gene expression profiles. This analysis led to the identication of three to seven different subtypes that were irrespective of the organ of origin. This analysis revealed that the majority of the subtype-specific gene signatures and mechanisms were conserved across different organs.

This study supports the clinical significance of targeting these tumor subtypes irrespective of the organ types. Future efforts are now needed to validate the MOT subtypes using clinical assays based on RT-PCR and/or immunohistochemistry needs. Subsequently, such methods could be further developed and tested for clinical utility. The current MOTassigner-275 and MOTassigner-51 signatures are a starting point to develop these assays and test their predictive values. Overall, our study provides the tools to test novel inhibitors without known pharmacodynamic markers (e.g. mutations) in selected patient population with similar subtype tumors.

## Methodology

### Selection of variable genes and merging of data sets

Gene expression (Affymetrix GeneChip® arrays) microarray data (CEL files) from different studies were obtained from Gene Expression Omnibus (GEO) (16), and they were preprocessed and normalized (robust multiarray average; rma(17)) using R and Bioconductor(18). GEOquery(19), a Bioconductor package was used to obtain the patient data from GEO data sets.

### NMF and Statistical Analysis of Microarrays (SAM) analysis

Datasets were subjected to NMF(15) based consensus clustering to identify subtypes. Later, the subtypes defined by NMF were used to perform SAM(20) analysis to identify genes that are specific to NMF identified subtypes. Both NMF and SAM analysis were described in our previous publication (3).

### Clustering of data and heatmaps

Gene Cluster 3.0(21) was used to cluster the gene expression profile data and the results were viewed using GenePattern-based Hierarchical Clustering Viewer(22).

## References

1. Alizadeh AA, et al. (2000) Distinct types of diffuse large B-cell lymphoma identified by gene expression profiling. Nature 403(6769):503-511.

2. Collisson EA, et al. (2011) Subtypes of pancreatic ductal adenocarcinoma and their differing responses to therapy. Nature medicine 17(4):500-503.

3. Sadanandam A, et al. (2013) A colorectal cancer classification system that associates cellular phenotype and responses to therapy. Nat Med: doi:10.1038/nm.3175.

4. Perou CM, et al. (2000) Molecular portraits of human breast tumours. Nature 406(6797):747-752.

5. TCGA (2012) Comprehensive molecular characterization of human colon and rectal cancer. Nature 487(7407):330-337.

6. Verhaak RG, et al. (2009) Integrated genomic analysis identifies clinically relevant subtypes of glioblastoma characterized by abnormalities in PDGFRA, IDH1, EGFR, and NF1. Cancer cell 17(1):98-110.

7. Lynch TJ, et al. (2004) Activating mutations in the epidermal growth factor receptor underlying responsiveness of non-small-cell lung cancer to gefitinib. The New England journal of medicine 350(21):2129-2139.

8. Slamon DJ, et al. (2001) Use of chemotherapy plus a monoclonal antibody against HER2 for metastatic breast cancer that overexpresses HER2. The New England journal of medicine 344(11):783-792.

9. Heiser LM, et al. (2011) Subtype and pathway specific responses to anticancer compounds in breast cancer. Proceedings of the National Academy of Sciences of the United States of America 109(8):2724-2729.

10. Chin K, et al. (2006) Genomic and transcriptional aberrations linked to breast cancer pathophysiologies. Cancer cell 10(6):529-541.

11. Jorissen RN, et al. (2009) Metastasis-Associated Gene Expression Changes Predict Poor Outcomes in Patients with Dukes Stage B and C Colorectal Cancer. Clin Cancer Res 15(24):7642-7651.

12. Bild AH, et al. (2006) Oncogenic pathway signatures in human cancers as a guide to targeted therapies. Nature 439(7074):353-357.

13. Badea L, Herlea V, Dima SO, Dumitrascu T, & Popescu I (2008) Combined gene expression analysis of whole-tissue and microdissected pancreatic ductal adenocarcinoma identifies genes specifically overexpressed in tumor epithelia. Hepato-gastroenterology 55(88):2016-2027.

14. Tothill RW, et al. (2008) Novel molecular subtypes of serous and endometrioid ovarian cancer linked to clinical outcome. Clin Cancer Res 14(16):5198-5208.

15. Brunet JP, Tamayo P, Golub TR, & Mesirov JP (2004) Metagenes and molecular pattern discovery using matrix factorization. Proceedings of the National Academy of Sciences of the United States of America 101(12):4164-4169.

16. Edgar R, Domrachev M, & Lash AE (2002) Gene Expression Omnibus: NCBI gene expression and hybridization array data repository. Nucleic acids research 30(1):207-210.

17. Irizarry RA, et al. (2003) Exploration, normalization, and summaries of high density oligonucleotide array probe level data. Biostatistics (Oxford, England) 4(2):249-264.

18. Gentleman RC, et al. (2004) Bioconductor: open software development for computational biology and bioinformatics. Genome biology 5(10):R80.

19. Sean D & Meltzer PS (2007) GEOquery: a bridge between the Gene Expression Omnibus (GEO) and BioConductor. Bioinformatics (Oxford, England) 23(14):1846-1847.

20. Tusher VG, Tibshirani R, & Chu G (2001) Significance analysis of microarrays applied to the ionizing radiation response. Proceedings of the National Academy of Sciences of the United States of America 98(9):5116-5121.

21. Eisen MB, Spellman PT, Brown PO, & Botstein D (1998) Cluster analysis and display of genome-wide expression patterns. Proceedings of the National Academy of Sciences of the United States of America 95(25):14863-14868.

22. Reich M, et al. (2006) GenePattern 2.0. Nature genetics 38(5):500-501.

